# Predicting heterogeneity in clone-specific therapeutic vulnerabilities using single-cell transcriptomic signatures

**DOI:** 10.1101/2020.11.23.389676

**Authors:** Chayaporn Suphavilai, Shumei Chia, Ankur Sharma, Lorna Tu, Rafael Peres Da Silva, Aanchal Mongia, Ramanuj DasGupta, Niranjan Nagarajan

## Abstract

While understanding heterogeneity in molecular signatures across patients underpins precision oncology, there is increasing appreciation for taking intra-tumor heterogeneity into account. Single-cell RNA-seq (scRNA-seq) technologies have facilitated investigations into the role of intra-tumor transcriptomic heterogeneity (ITTH) in tumor biology and evolution, but their application to *in silico* models of drug response has not been explored. Based on large-scale analysis of cancer omics datasets, we highlight the utility of ITTH for predicting clinical outcomes. We then show that heterogeneous gene expression signatures obtained from scRNA-seq data can be accurately analyzed (80%) in a recommender system framework (CaDRReS-Sc) for *in silico* drug response prediction. Patient-derived cell lines capturing transcriptomic heterogeneity from primary and metastatic tumors were used as *in vitro* proxies for validating monotherapy predictions (Pearson r>0.6), as well as optimal drug combinations to target different subclonal populations (>10% improvement). Applying CaDRReS-Sc to the increasing number of publicly available tumor scRNA-seq datasets can serve as an *in silico* screen for further *in vitro* and *in vivo* drug repurposing studies.

**Graphical abstract:** 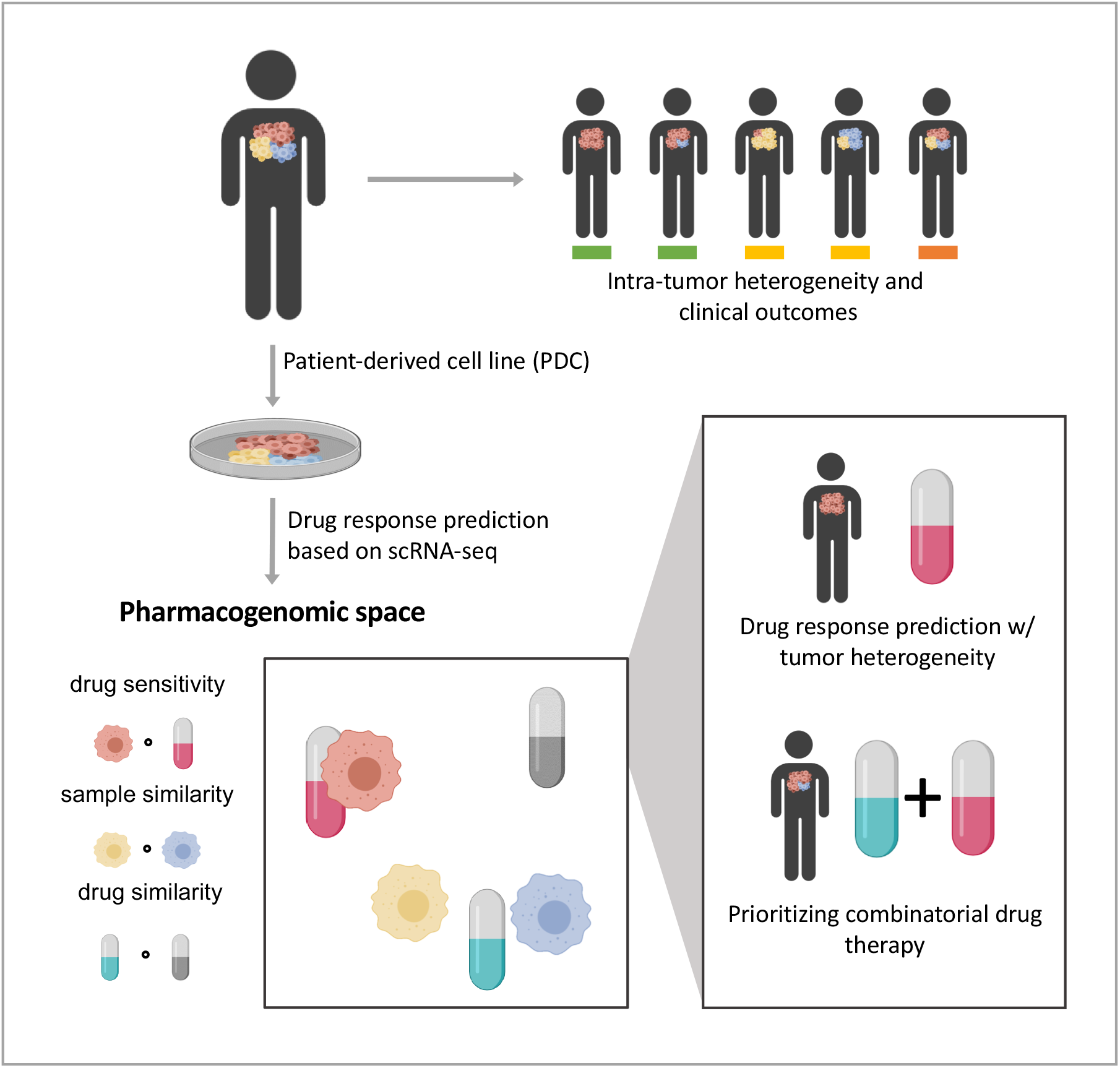

**Highlights:**

- Large-scale analysis to establish the impact of transcriptomic heterogeneity within tumors on clinical outcomes
- Calibrated recommender system for drug response prediction based on single-cell RNA-seq data (CaDRReS-Sc)
- Prediction of drug response in patient-derived cell lines with transcriptomic heterogeneity
- *In silico* identification of drug combinations that work based on clonal vulnerabilities

## Introduction

Tumors comprise of heterogeneous populations of malignant cells that display cellular plasticity and phenotypic heterogeneity, as determined by genetic and environmental cues (Lee et al., 2018; McGranahan and Swanton, 2017; Meacham and Morrison, 2013). Phenotypic heterogeneity in cancer cells is defined by transcriptomic signatures that govern cell biological behaviors, such as proliferation, apoptosis, migration, invasion, metabolism and immune response (Neftel et al., 2019; Puram et al., 2017). Intra-tumor transcriptomic heterogeneity (ITTH) can confer differential selective advantages to influence tumor progression and metastasis *in vivo* (Bruna et al., 2016; Wei et al., 2016), as well drug response *in vitro* (Chia et al., 2017; Sharma et al., 2018).

Advances in high-throughput sequencing have enabled large-scale studies into inter-patient tumor heterogeneity at the molecular level (Ding et al., 2018; Hoadley et al., 2018; Zhang et al., 2011), serving as the basis to distinguish cancer subtypes, investigate tumor biology and define treatment regimens (Lee et al., 2018; Schilsky, 2010). These efforts have been complemented by studies on cancer cell lines (Barretina et al., 2012; Iorio et al., 2016; Rees et al., 2016) to understand the relationship between molecular markers and drug response *in vitro*. Several machine learning models have been proposed to utilize information from multi-omic profiles to predict drug response for cell lines (Ammad-ud-din et al., 2016; Azuaje, 2016; Baptista et al., 2020; Basu et al., 2018; Suphavilai et al., 2018; Wang et al., 2017), although significant challenges remain in terms of robustness, generalizability and translatability into the clinic. In particular, existing models do not explicitly account for intra-tumor transcriptomic heterogeneity and have primarily been trained and tested on clonal cell lines.

In this work, we begin by highlighting the impact of intra-tumor transcriptomic heterogeneity on clinical outcomes based on large-scale re-analysis of TCGA data (Ding et al., 2016; Liu et al., 2018). We then develop a framework that robustly combines single-cell RNA-sequencing (scRNA-seq) data with a recommender system (Suphavilai et al., 2018) to predict drug response heterogeneity in a tumor (CaDDReS-Sc). ScRNA-seq data from 12 patient-derived cell lines (PDCs) and cell viability measurements in response to 8 drugs under 2 doses was used to evaluate robustness of CaDDReS-Sc predictions to intra-tumor transcriptomic heterogeneity. Extending to combinations of drugs, we show that drug pairs identified *in silico* by CaDDReS-Sc to optimally inhibit transcriptionally distinct cell clusters were more effective than individual drugs *in vitro*.

## Results

### Intra-tumor transcriptomic heterogeneity is significantly associated with treatment response and patient outcomes

To investigate the relationship between intra-tumor transcriptomic heterogeneity (ITTH) and clinical outcomes, we leveraged transcriptomic data from TCGA for 10,956 tumors across 33 cancer types, and an *in silico* deconvolution approach (Newman et al., 2015), to define a transcriptomic heterogeneity score for each patient (ITTH score, measuring degree of heterogeneity in gene expression across cells of a tumor inferred based on bulk transcriptomic profiles; **Methods**). Comparing these *in silico* heterogeneity scores with single-cell RNA-seq derived gold-standards (scITTH score; **Methods**) on two different datasets (Puram et al., 2017; Sharma et al., 2018) showed that the *in silico* scores provided a useful proxy to capture transcriptomic heterogeneity (Pearson r≥0.55; **Figure 1A**).

**Figure 1:**
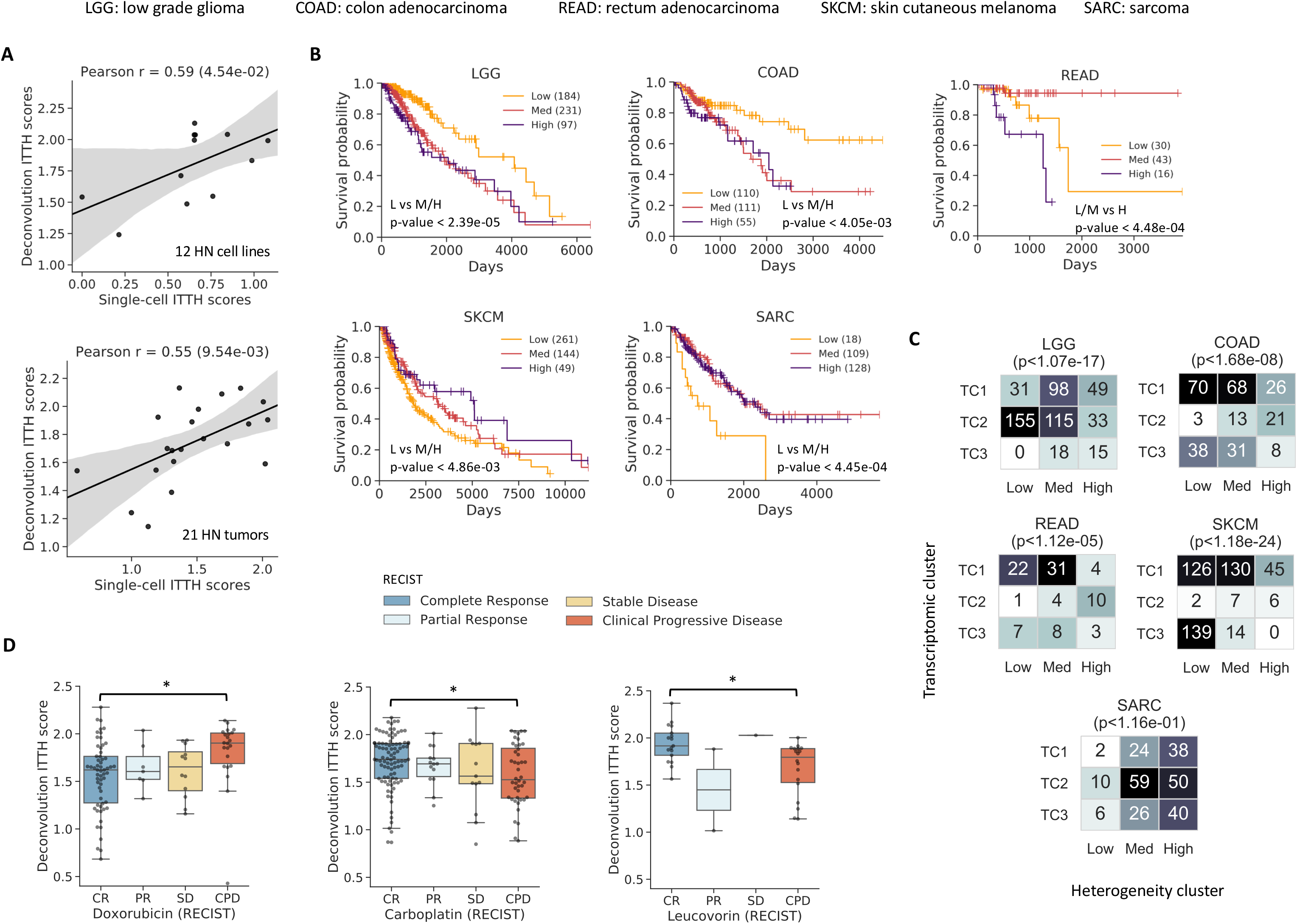
Impact of intra-patient transcriptomic heterogeneity on clinical outcomes. (A) Scatterplots showing the correlation between transcriptomic heterogeneity estimates based on *in silico* deconvolution (ITTH score) versus single-cell analysis derived values (scITTH score). (B) Survival analysis with ITTH clusters (Low/Medium/High) identified significant differences across various cancer types (FDR-corrected log-rank p-value<0.05). (C) Plots depicting the overlap between clusters based on transcriptomic profiles (TC) and ITTH scores. P-values are based on Fisher’s exact test and indicate that the clusters are distinct for most cancer types. (D) Comparison between ITTH scores of patients from different RECIST classes for Doxorubicin, Carboplatin and Leucovorin highlighting significant differences (*: FDR-corrected Wilcoxon p-value<0.05).

Survival data for various cancer types was then analyzed to detect differences in patients with low, medium, and high transcriptomic heterogeneity (24/33 cancer types with ≥15 samples in each class). Significant associations between ITTH and survival were observed in 5 cancer types, with high heterogeneity associated with poorer outcomes in some cancer types and low heterogeneity in others (**Figure 1B**). The most significant associations were observed in low-grade glioma (LGG, Low vs Med/High; FDR-corrected log-rank p-value<2.39×10^−5^) and sarcoma (SARC, Low vs Med/High, p-value<4.45×10^−4^), in agreement with prior work on the impact of cell-type diversity in low-grade glioma (Easwaran et al., 2014) and cellular plasticity in sarcoma (Suvà et al., 2009) on treatment outcomes.

To investigate if information in ITTH clusters is captured directly in clustering based on bulk transcriptomic profiles, corresponding clusters were compared for the 5 cancer types (LGG, SARC, COAD, READ, SKCM; **Figure 1C**; **Methods**). Among these 5 cancer types, associations were observed between transcriptomic clusters and survival in 2 cancer types (LGG, SKCM; FDR-corrected log-rank p-value<0.05) and these transcriptomic clusters were typically observed to be orthogonal to ITTH clusters (**Figure 1C**; 4/5 cancer types; Chi-squared test p-value<0.05). For example, in low-grade glioma, the low ITTH cluster is characterized by better survival rate compared to transcriptomic clusters 1 and 2 (TC1, TC2), while in rectum adenocarcinoma, the high ITTH cluster is characterized by lower survival rate compared to all three transcriptomic groups (**Supplementary figure 1**), highlighting the additional information captured in ITTH analysis.

Drawing on the availability of patient response data in TCGA for a few drugs (n=8) and cancer types (n=24), we systematically assessed associations between ITTH scores and clinical drug response (**Figure 1D**, **Supplementary figure 2**). Significant associations were identified in 3/8 drugs (Doxorubicin, Carboplatin, Leucovorin; FDR-corrected Wilcoxon p-value<0.05; **Figure 1D**; **Methods**), where for example, Doxorubicin-resistant patients exhibited significantly higher transcriptomic heterogeneity (n=80; CR vs CPD Wilcoxon p-value< 9.86×10^−3^). This response pattern for Doxorubicin in patients with high ITTH score could indicate pre-existing resistant populations (Dagogo-Jack and Shaw, 2018), tumor evolution (Sharma et al., 2018) or high variability in drug-target engagement (Sparks et al., 2018). For Carboplatin the opposite trend was observed (**Figure 1D**; n=138; CR vs CPD Wilcoxon p-value<2.06×10^−2^), where responders showed significantly higher transcriptomic heterogeneity, consistent with prior work (Böttger et al., 2019). Direct measurement and incorporation of transcriptomic heterogeneity could therefore lead to more accurate predictions for drug response, as we explore in the next section.

**Figure 2:**
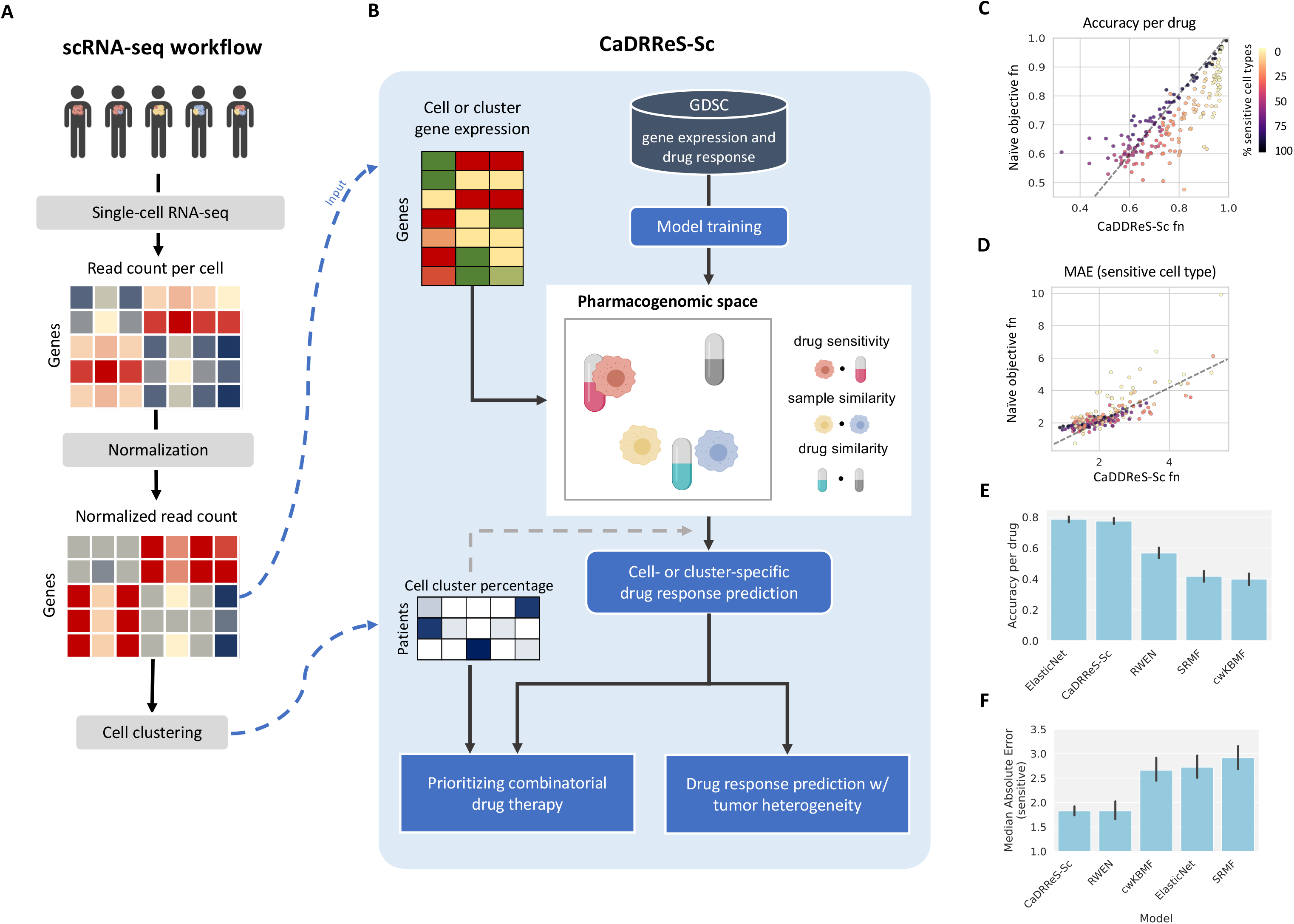
CaDRReS-Sc accurately predicts drug response in unseen cell types. (A) Overview of single-cell RNA-seq workflow to preprocess sequencing data and provide inputs to CaDRReS-Sc (indicated by blue dashed lines). (B) Overview of CaDRReS-Sc workflow, where a pre-trained *pharmacogenomic space* based on drug response and gene expression profiles from cell-line experiments is used to provide cell- or cluster-specific drug responses predictions. These are then combined to estimate overall drug response and prioritize drug combinations for a patient. (C) Comparison of prediction accuracy on unseen cell types between CaDRReS-Sc’s objective function and a naïve function that does not take uncertainty in IC50 values into account. Each dot represents a drug (n=226) and dot colors represent the percentage of sensitive cell lines. As can be seen here, CaDRReS-Sc’s objective function is particularly useful when the percentage of sensitive cell lines is low. (D) Comparison of median absolute error (MAE) obtained based on predictions using CaDRReS-Sc as well as a naïve objective function. CaDRReS-Sc’s robust objective function results in lower MAE across a majority of drugs (points above the y=x line), especially for drugs with a lower percentage of sensitive cell lines (lighter shades). (E) Histograms showing the average prediction accuracy (error bars show 1 standard deviation) using different drug response prediction approaches. (F) Histograms showing MAE (error bars show 1 standard deviation) with different drug response prediction approaches. Overall, CaDRReS-Sc was seen to have high accuracy on the sensitive/non-sensitive classification task while reporting the lowest MAE for the IC50 regression task.

### Calibrating a recommender system for improved predictive performance on diverse unseen cell types

The development of single-cell transcriptomics has enabled direct identification and quantification of cell populations within a tumor (Kim et al., 2015; Patel et al., 2014; Puram et al., 2017; Sharma et al., 2020). Corresponding scRNA-seq workflows with gene expression measurement, normalization, cell clustering and summarization (**Figure 2A**), can be coupled in principle with existing methods that predict drug response from bulk transcriptomic profiles (Ammad-ud-din et al., 2016; Basu et al., 2018; Iorio et al., 2016; Suphavilai et al., 2018; Wang et al., 2017) to obtain cell-specific response information. While the utility of such a workflow and potential techniques to obtain a summarized response score for the tumor have not been explored, a more fundamental challenge is the robustness of such models to diverse, unseen cell types (Baptista et al., 2020).

To address these questions, we extended an existing recommender system trained with cancer cell line data (Suphavilai et al., 2018) for improved robustness on diverse, unseen cell types, and the ability to combine cell-specific predictions into accurate tumor response values (CaDRReS-Sc; **Figure 2B**). Specifically, we designed a novel objective function that enables the model to simultaneously classify sensitive/resistant cell types and predict half-maximal inhibitory concentration (IC50) values for sensitive cases (**Supplementary figure 3**; **Methods**). Comparison of predictive accuracy versus a naïve objective function (mean squared error for IC50) on unseen cell lines showed significant improvements (**Figure 2C**; 5-fold cross-validation; Wilcoxon p-value<2.66×10^−4^), especially for drugs with a smaller proportion of sensitive cell lines. By focusing on predicting response values for sensitive cell lines, we observed that the overall median absolute error (MAE) was reduced in a majority of the drugs (**Figure 2D**; 5-fold cross-validation; Wilcoxon p-value<1.80×10^−5^; **Methods**), enabling accurate prediction of drug response at specific dosages.

**Figure 3.**
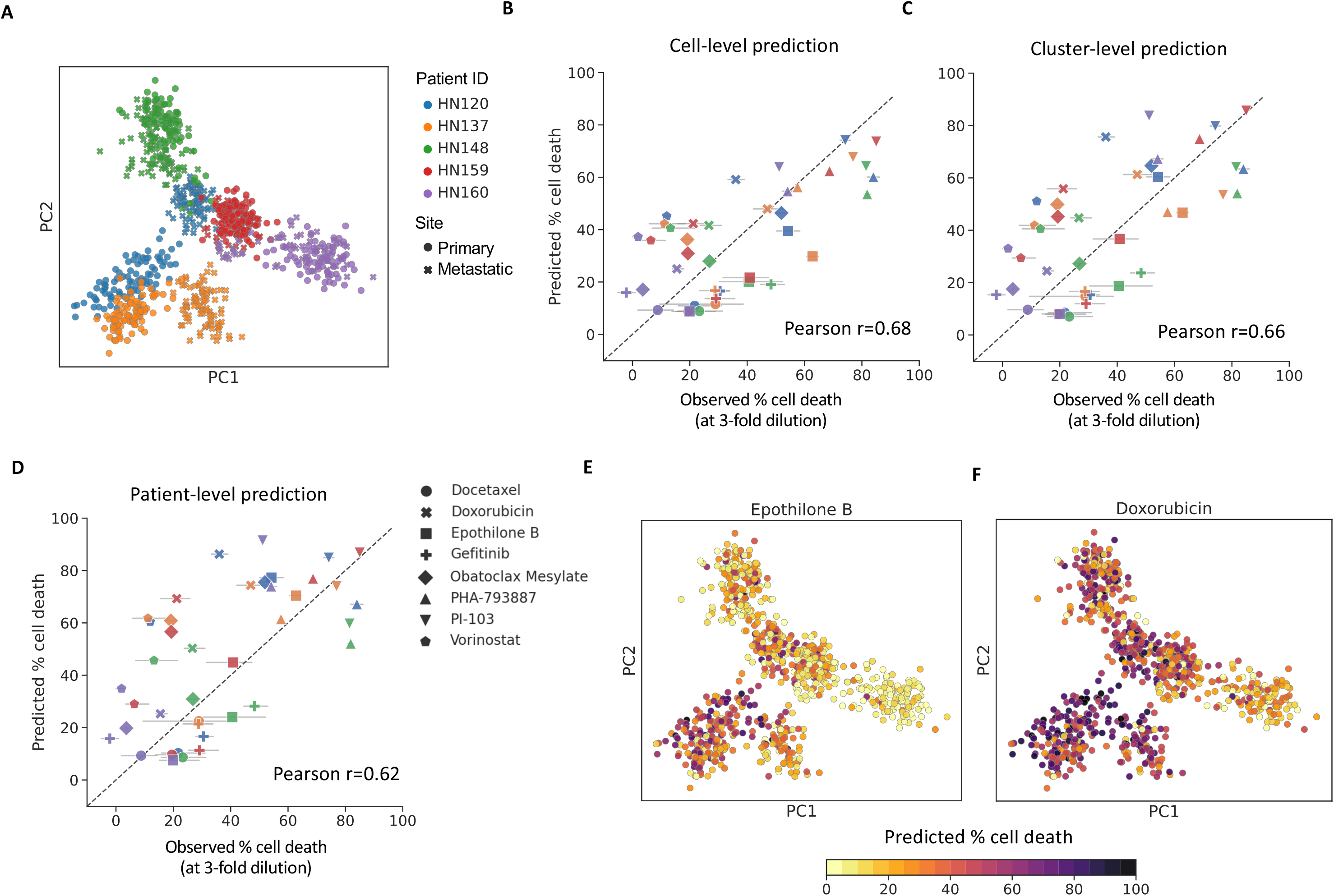
Calibrated drug response prediction in heterogenous patient-derived cell lines using scRNA-seq data. (A) PCA plot showing the diversity of single-cell transcriptomic profiles from different patient-derived cell lines. Comparison of observed and predicted cell death percentages for 5 patient-derived cell lines using 8 different drugs (at lower concentrations), based on CaDRReS-Sc analysis at the (B) cell-level, (C) cluster-level and (D) patient-level. Error bars show 1 standard deviation based on 3 experimental replicates. Note that that cell and cluster-level predictions show greater correlation with experimental observations than patient-level predictions, highlighting the utility of scRNA-seq data. (E-F) PCA plots showing varied cell-level response predictions to treatment with Epothilone B and Doxorubicin, highlighting substantial inter- and intra-patient drug response heterogeneity.

Benchmarking against other machine learning based approaches for drug response prediction trained on the same cancer line dataset, such as ElasticNet (Iorio et al., 2016), cwKBMF (Ammad-ud-din et al., 2016), SRMF (Wang et al., 2017) and RWEN (Basu et al., 2018), we noted that average prediction accuracy for CaDRReS-Sc and ElasticNet was significantly better than other methods (cwKBMF, SRMF, RWEN; Wilcoxon p-value<0.05), with an average prediction accuracy of around 80% compared to <60% for other methods (**Figure 2E**). This improvement seems to be due to higher accuracy for drugs with a smaller fraction of sensitive cell lines, where CaDRReS-Sc is able to recapitulate the performance of a drug-specific regression model (ElasticNet) while learning a shared model across drugs (similar to cwKBMF, SRMF and RWEN; **Supplementary figure 4A**). In addition, we observed that by aggregating information across drugs, CaDRReS-Sc is able to predict more precise response values across different dosages (**Supplementary figure 4B**), reducing MAE by >30% compared to cwKBMF, ElasticNet and SRMF (**Figure 2F**). Finally, we confirmed that a numerical integration based approach to combine drug response values across cell clusters, accurately predicts overall tumor response (**Supplementary figure 5**). Together, these capabilities enable CaDRReS-Sc to accurately predict drug response in the presence of transcriptomic heterogeneity as evaluated in the next section based on scRNA-seq data from patient-derived cell lines.

**Figure 4.**
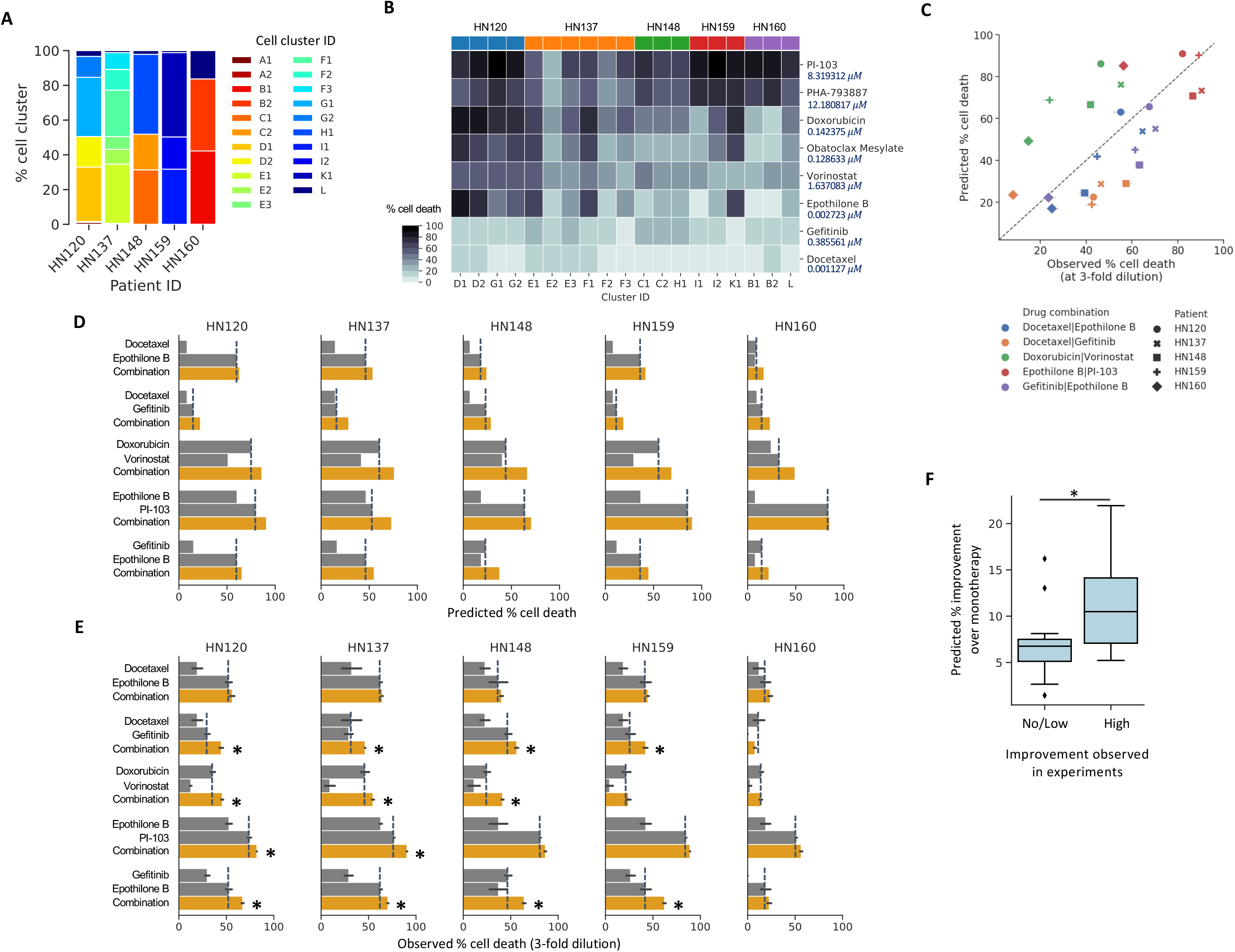
Prioritizing drug combinations targeting transcriptionally-distinct subclones with CaDRReS-Sc. (A) Proportions of various transcriptionally-distinct cell clusters (n=21) in head and neck cancer patient-derived cell lines. (B) Heatmap of predicted cell death percentages across cell clusters within each patient. (C) Comparison between predicted and observed drug response for five different drug combinations and patient-derived cells. Boxplots contrasting monotherapy (grey) and combinatorial therapy (orange) response based on, (D) CaDRReS-Sc predictions and (E) experimental measurements. Error bars show 1 standard deviation (n=2-3), dashed lines indicate the best monotherapy, and asterisk symbols indicate drug combinations that show improvement. In general, relative response values for monotherapy and combinatorial therapy, as observed from experimental measurements, were also reflected in CaDRReS-Sc predictions. (F) Boxplots showing that drug combinations that were observed to improve over monotherapy (x-axis, no/low vs high determined based on median value in experiment) had significantly higher predicted improvements (combination over monotherapy) using CaDRReS-Sc as well (y-axis).

### Accurate drug response prediction in the presence of intra-patient heterogeneity

As commonly used cancer cell lines typically lack significant transcriptomic heterogeneity, we leveraged patient-derived cell lines (PDCs) to serve as model systems where sensitivity measurements can be systematically and conveniently made across multiple drugs, while capturing *in vivo* transcriptomic heterogeneity (Chia et al., 2017). In total, 12 PDCs from head and neck cancer patients (Sharma et al., 2018) were used for scRNA-seq analysis (median >10^5^ reads/cell, >5×10^3^ detected genes, >1,200 cells in total; **Supplementary figure 6**; **Methods**) and drug response was measured for 8 different drugs at 2 different concentrations (median IC50 of ATCC head and neck cancer cell lines and 3-fold lower; **Methods**). Visualization of single-cell transcriptomic profiles in 2D space confirmed that significant intra-patient transcriptomic heterogeneity was seen in PDCs (relative to inter-patient heterogeneity; **Figure 3A**), particularly when primary and lymph-node metastatic tumors are considered (e.g. HN120 and HN137).

We explored several strategies to utilize scRNA-seq data — ranging from using transcriptomic profiles of individual cells, aggregating profiles within a cluster of cells, to combining profiles at the patient-level — for predicting drug response (cell death percentage at a specific drug dosage; **Methods**). Comparing predicted and observed drug response (5 pooled PDCs; 8 drugs) we observed significant correlations using CaDRReS-Sc under all three strategies (**Figure 3B-D**; Pearson r=0.68, 0.66, 0.62, p-value<1.11×10^−6^, 3.59×10^−6^, 1.93×10^−5^, respectively). Despite noise and dropout events observed in single-cell data (Kolodziejczyk et al., 2015), predictions based on cell- and cluster-level transcriptomic profiles consistently showed better agreement with *in vitro* drug response compared to patient-level prediction (Pearson r=0.68/0.66 vs 0.62; consistently across drug dosages, **Supplementary figure 7**), highlighting the importance of transcriptomic heterogeneity and the robustness of kernel-based predictions with CaDRReS-Sc (Pearson r≤0.57 with ElasticNet and RWEN at cell-level).

As CaDRReS-Sc is based on a *pharmacogenomic space* model that can help interrogate drug-response mechanisms (Suphavilai et al., 2018), we applied it to our single-cell data to study drug-pathway associations for individual cells (**Methods**). For example, we found a wide range of responses for Epothilone B (**Figure 3E**), especially amongst cells in HN120 and HN137 where primary cells are more sensitive than metastatic cells (**Supplementary figure 8A**). Examination of CaDRReS-Sc’s latent pharmacogenomic space identified a significant association between Wnt pathway activation and Epothilone B response (Wilcoxon p-value<7.24×10^−8^; **Supplementary figure 8C**), consistent with prior work on this subject (Ciani et al., 2004). Similarly, we noted diverse responses across cells for Doxorubicin (**Figure 3F**; e.g. primary cells tend to be more sensitive in HN120, **Supplementary figure 8B**), and significant association with activation of the Fas pathway (Wilcoxon p-value<4.69×10^−15^; **Supplementary figure 8D**) (Kim et al., 2009), highlighting the potential to obtain biological insights on therapeutic vulnerabilities based on single-cell information and the interpretability of the CaDRReS-Sc model.

### Drug combinations can be identified *in silico* by utilizing scRNA-seq data

Going beyond monotherapy, the ability to predict drug combinations to target different cancer cell types within a heterogeneous tumor can be essential for improving treatment efficacy in the clinic (Dancey and Chen, 2006; Jia et al., 2009). The utility of combinations can arise from independent drug action as well as the increased chance of specific clones being sensitive to a drug (Palmer and Sorger, 2017). To explore this, we evaluated if *in silico* predictions with scRNA-seq data could reveal drug combinations that provide better response compared to monotherapy, even at lower dosages. Specifically, by inspecting 21 cell clusters across all PDCs identified in our monotherapy analysis, we observed different cluster proportions across patients (**Figure 4A**) and a broad range of predicted monotherapy responses across cell clusters (**Figure 4B**). These results suggest variability in therapeutic response across different subclones in a given individual, allowing us to identify complementary drug combinations for subclones.

We calculated an expected combination effect (% cell death) for five candidate drug combinations, including Docetaxel:Epothilone B, Docetaxel:Gefitinib, Gefitinib:Epothilone B, Epothilone B:PI-103, and Doxorubicin:Vorinostat, based on predicted cluster-specific drug responses and the distribution of cell clusters. These combinations were then functionally validated on five pooled PDCs via cell-based viability assays using low drug dosages to circumvent off-target effects resulting from extreme inhibition (**Methods**). Despite the potential for drug interactions (Scripture and Figg, 2006), CaDRReS-Sc predictions for response to various drug combinations showed a clear correlation to observed responses across the 25 different experimental conditions (**Figure 4C**; Pearson r=0.58; p-value<2.30×10^−3^), in comparison to weaker correlations with other methods (Pearson r≤0.49 with ElasticNet and RWEN).

Testing drugs in each combination at low dosages also helped to mimic what might be needed to support the mitigation of side effects from combinatorial treatment (Jia et al., 2009). We then evaluated if this approach can be used to predict pairs of drugs that can elicit greater overall cell death compared to monotherapy. Overall, we observed consistent trends between *in silico* predictions with CaDRReS-Sc (**Figure 4D**) and experimental results (**Figure 4E**). For instance, the combination of Doxorubicin and Vorinostat was predicted to provide a notable improvement over monotherapy (+22%) in HN148, which was observed experimentally as well (+11%), consistent with prior work on this combination (Cheriyath et al., 2011). By computing the expected improvement of combinatorial therapy over monotherapy, we observed concordance between CaDRReS-Sc’s *in silico* predictions and *in vitro* experimental results (**Figure 4F**; 25 different experimental conditions; Wilcoxon p-value<3.39×10^−2^), but no significant associations for other methods (ElasticNet, RWEN). These results indicate that CaDRReS-Sc can sufficiently capture therapeutic response for mono- and combinatorial therapy, enabling prioritization of drugs and combinations for *in vitro* and *in vivo* studies.

## Discussion

While the role of intra-patient heterogeneity in genetic mutations has been extensively explored with respect to tumor biology (Bhatia et al., 2012; Meacham and Morrison, 2013), fewer studies have investigated how this combines with epigenetic heterogeneity to influence transcriptomic heterogeneity (Easwaran et al., 2014), drug response and patient outcomes (Lee et al., 2018; McGranahan and Swanton, 2017). In this work, we leveraged the availability of large-scale tumor sequencing datasets to highlight the relationship between intra-tumor transcriptomic heterogeneity and patient outcomes, identifying associations in 5 out of 24 cancer types and 3 out of 8 standard-of-care drugs. While this analysis emphasized the general importance of taking ITTH into account for predicting treatment response and outcomes, the power to detect associations might have been limited by the dependence on an *in silico* deconvolution approach (Gentles et al., 2015; Newman et al., 2015). With the increasing availability of single-cell tumor sequencing datasets, the resolution of such analysis could be greatly improved and help identify shared cell populations that contribute to treatment resistance across patients.

Predicting treatment response *in silico* in the presence of intra-tumor heterogeneity requires models that provide calibrated values for a single drug across many cell types, while prior work has focused on calibrated predictions for a cell type across many drugs (Suphavilai et al., 2018). To address this, CaDRReS-Sc uses a novel objective function that accounts for the uncertainty in drug response values across drugs. This allowed CaDRReS-Sc to train a model that is as accurate as single-drug models (80%), while leveraging information across drugs to provide highly calibrated response values compared to start-of-the-art methods. Furthermore, CaDRReS-Sc’s latent pharmacogenomic model provides ready visualization and interpretation to examine the pathways involved in drug response heterogeneity in a tumor.

Patient-derived cell lines (PDCs) serve as ideal systems for drug sensitivity measurements *in vitro* while capturing intra-tumor transcriptomic heterogeneity (Chia et al., 2017; Sharma et al., 2018), and we leveraged this in a proof-of-concept study, with 12 head and neck cancer PDCs and 8 drugs under 2 dosages, to assess the ability to predict drug response *in silico* in the presence of transcriptomic heterogeneity. Despite variations in experimental conditions between training data from public cancer cell line datasets (Iorio et al., 2016) and test response data from heterogenous PDCs, *in silico* predictions from CaDRReS-Sc could recapitulate cell death percentages observed in our *in vitro* experiments (Pearson r=0.68, **Figure 3B**), highlighting the robustness of such models. Further availability of drug response data in PDCs and at clinically relevant doses (Liston and Davis, 2017), could help advance predictive performance and clinical utility of such models.

In predicting response to monotherapies, we observed consistently higher correlations with *in vitro* measurements when using transcriptomic profiles with higher granularity (individual cells or cell clusters versus bulk profiles). This prompted us to consider prioritizing combinatorial therapy options based on CaDRReS-Sc predictions for different subclones assuming that the combinatorial effect can be approximated in many cases through independent drug action on distinct subclonal cancer cell populations (Palmer and Sorger, 2017). Although this does not directly account for the impact of drug-drug interactions (Chou, 2010; Jia et al., 2009; Yadav et al., 2015), overall we were able to capture the effect of combinatorial drug therapy (**Figure 4C-E**) and its improvement over monotherapy (**Figure 4F**). We envisage, therefore, that the growing corpus of scRNA-seq data can be data-mined using CaDRReS-Sc to identify drug combinations that target clone-specific therapeutic vulnerabilities and lead to better treatment outcomes (Dagogo-Jack and Shaw, 2018; Sharma et al., 2018).

Developing *in silico* tools for predicting *in vivo* treatment response remains a challenge as multiple factors (e.g. tumor microenvironment, immune response, overall patient health) can impact patient trajectories. In this study, we aimed to bridge the *in silico* to *in vitro* gap for predicting response to mono- and combinatorial therapy in the presence of transcriptomic heterogeneity. Together with improved technologies for patient-derived cancer-cell models, this combined *in silico*/*in vitro* approach could form the basis of a first-cut precision oncology platform that prioritizes mono- and combinatorial therapy options in a clinically-relevant timeframe.

## Acknowledgements

This work was supported by funding from A*STAR.

## Author Contributions

C.S., N.N., R.D., A.S. and S.C. planned and designed the project. C.S. developed CaDRReS-Sc models and performed all computational analysis with N.N.’s supervision. S.C. and A.S. planned wet-lab experiments and S.C. conducted them with R.D.’s supervision. L.T. performed ITTH analysis in the TCGA cohort with C.S. and N.N.’s supervision. R.P. and A.M. performed additional CaDRReS-Sc analysis and developed documentation with C.S.’s guidance. C.S., S.C., R.D. and N.N. wrote the manuscript with input from all authors.

## Code Availability

A Python package for CaDRReS-Sc and example scripts for predicting drug response based on scRNA-seq data are available at https://github.com/CSB5/CaDRReS-Sc. With scRNA-seq data, users can calculate aggregated gene expression profiles for clusters, predict cell-specific or cluster-specific drug responses using a pre-trained model, and estimate overall drug response for a patient.

## Methods

### Datasets and preprocessing

#### Tumor data

Gene expression (FPKM-UQ normalized RNA-seq) and patient survival data for 10,956 tumors from 33 cancer types were obtained from The Cancer Genome Atlas (TCGA) Research Network (https://www.cancer.gov/tcga). Clinical drug response information and *Response Evaluation Criteria in Solid Tumors* (RECIST) values were obtained from prior work to curate drug response information (Ding et al., 2016), and statistical analysis was limited to drugs with sufficient number of patients with clinical data (n=8 with ≥15 patients in “Complete Response” and “Clinical Progressive Disease” classes).

#### Cancer cell line data for model training

Drug response data and RMA-normalized gene expression data for 1,074 cancer cell lines and 226 drugs tested at nine different dosages were obtained from the GDSC database (Iorio et al., 2016) to be used for model training. For each gene, log2 expression fold-change was calculated with respect to its average expression across cell lines, and cell line kernel features were calculated using Pearson correlation based on 1,856 essential genes (Suphavilai et al., 2018; Wang et al., 2015). Models were trained to predict half-maximal inhibitory concentrations (IC50) based on Bayesian sigmoid curve fitting estimates (Suphavilai et al., 2018).

#### Single-cell RNA-seq data

Single-cell RNA-seq data for 1,241 cells from 12 head and neck patient-derived cell lines was obtained based on a previously published study (Sharma et al., 2018). Read counts per gene were obtained by mapping reads with STAR (v2.5.2a, default parameters), followed by RSEM analysis (v1.3.0, default parameters). Cells with <10,000 reads and a cell with a large number of expressed genes (n=14,558) were removed (the median number of genes per cell is 7,379; **Supplementary figure 6**). Genes expressed in <5% of cells were then filtered out to obtain expression values for 15,144 genes from 1,171 cells that were used for further analysis (as TPM values; **Supplementary file 1**). To evaluate ITTH scores, an additional scRNA-seq dataset containing 5,902 cells (TPM values for 23,686 genes) from 18 head and neck cancer patients was also obtained (Puram et al., 2017).

### Single-cell clustering and cluster-specific transcriptomic profiles

A standard scRNA-seq workflow described in the Scanpy tutorial was used to perform single-cell clustering (Butler et al., 2018; Wolf et al., 2018). Starting from the TPM matrix, cells with a large proportion (25%) of mitochondrial genes were removed as the high proportions are indicative of poor-quality cells (Lun et al., 2016). Expression values were log-normalized and adjusted based on detection of highly-variable-genes. The neighborhood graph was generated with n_neighbors=10 and n_pcs=40. In the final step, the neighborhood graph was used for cell clustering using the Louvain algorithm (Blondel et al., 2008).

To obtain a higher resolution of clustering, subclusters of large clusters (≥ 50 cells) were identified using the same process as the first round clustering. In total, 23 clusters were identified for the Sharma *et al* dataset (1,171 cells) and 62 clusters for the Puram *et al* dataset (5,902 cells) (**Supplementary file 2**). Transcriptomic profiles for each cell cluster (patient) were obtained by averaging TPM values across cells, to be used later for cluster-level (patient-level) drug response prediction.

### *In silico* deconvolution and intra-tumor transcriptomic heterogeneity

Percentages of pre-defined cancer cell types in each tumor were identified by using CIBERSORT (Newman et al., 2015), a tool for tumor deconvolution based on transcriptomic information. Since CIBERSORT requires a cell signature matrix containing gene expression profiles of specific cell types, we followed CIBERSORT’s manual to construct a new signature matrix using GDSC histological subtypes (n=53) to obtain a signature matrix with 1,529 marker genes.

To measure the degree of heterogeneity for each sample based on the deconvolution result, we defined an intra-tumor transcriptomic heterogeneity score (ITTH) as information entropy of the corresponding cell type profile i.e. *ITTH* = − Σ_*i*_ *P*_*i*_*logP*_*i*_, where *P*_*i*_ is the fraction of cells with cell type *i* identified in the tumor (**Supplementary file 3**). Cell types with <5% frequency were excluded to reduce the impact of classification noise and obtain a robust score. For patients with multiple samples in the TCGA dataset, an average ITTH score was used. Patients were classified at the pancan-level into three categories (low, medium, and high) based on the first and third quartiles of ITTH scores. Gene expression profiles for tumors from TCGA were clustered using non-negative matrix factorization (NMF; k = 3 to mimic the number of ITTH clusters) (Berry et al., 2007).

For scRNA-seq data, a proxy for bulk gene expression values was obtained by calculating the average gene expression across all cells and used to computed ITTH scores as described above. Gold-standard ITTH scores (single-cell ITTH) were then computed based on single-cell clustering (as described above) and then computing information entropy as before i.e. − Σ_*j*_*P*_*j*_*logP*_*j*_, where *P*_*j*_ is the fraction of cells that belong to cluster *j*.

### The CaDRReS-Sc framework

#### Learning a pharmacogenomic space

A pharmacogenomic space is a latent space that captures the relationship between drugs and samples (transcriptomic profiles for cells, cell clusters, cell lines, or patients), where a dot product between drug and sample vectors captures drug sensitivity. The pharmacogenomic space is learned in CaDRReS-Sc based on both transcriptomic and drug response profiles across multiple samples and drugs. The original objective function proposed in (Suphavilai et al., 2018) was defined as follow:

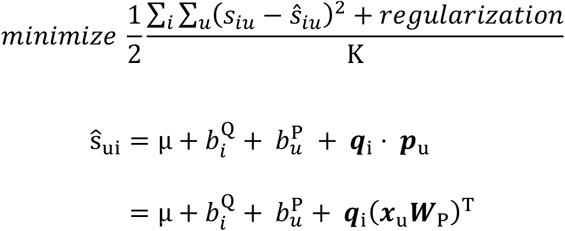

where *s*_*iu*_, the observed sensitivity score of sample *u* to drug *i*, is defined by *s*_*iu*_ = −log_2_(*IC*_50_), 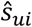 is the predicted sensitivity score, K is the total number of drug-sample pairs, *μ* is the overall mean drug response, 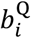 and 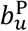 are bias terms for drug *i* and sample *u*, vectors ***q***_*i*_, ***p***_*u*_ ∈ ℝ^*f*^ represent drug *i* and sample *u* in the *f*-dimensional latent space, and ***W***_*P*_ ∈ ℝ^*d*^ ×^*f*^ is a transformation matrix that projects transcriptomic kernel features ***x***_*u*_ ∈ ℝ^*d*^ for each sample onto the pharmacogenomic space.

As estimates of 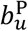 do not accurately capture the true bias of an unseen sample, the bias terms μ and 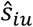 were removed from CaDRReS-Sc’s objective function, allowing sample bias to be implicitly captured in ***p***_u_. Furthermore, to reduce noise from extrapolation errors for IC50 values (**Supplementary figure 3**), a logistic weight function was introduced to assign a weight for each sample-drug pair as:

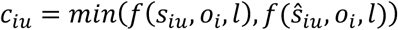

where *f* is a logistic function with slope *l* centered at *o*_*i*_, which is the maximum testing dosage for drug *i*. In resistant cases, the dose-response curve is extrapolated and *IC*_50_ estimates are higher than the maximum tested dosage. As a result, if both predicted 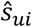 and observed *s*_*iu*_ dosages are greater than the maximum dosage, then *c*_*iu*_ is close to 0 and the error relative to the extrapolated *IC*_50_ value is down-weighted. Finally, to obtain an cancer type-specific model, *d*_*u*_ > 1 was defined as a weight of training sample *u* from a given cancer type, enabling the model to focus on accuracy for a subset of training samples. The final objective function used for learning the pharmacogenomic space was defined as:

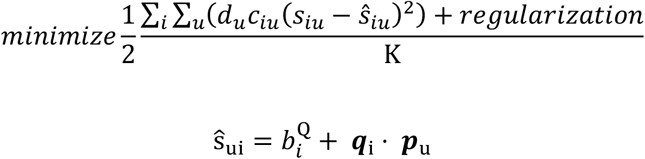

#### Model training and evaluation

The CaDRReS-Sc model was trained with a 10-dimensional pharmacogenomic space (*f* = 10), learning rate of 0.01, and number of epochs set to 100,000. Performance on unseen samples was estimated with 5-fold cross-validation within the GDSC dataset, and predictive performance for each drug was measured in terms of prediction accuracy and median absolute error (MAE). Drug-sample pairs were classified into two classes based on their IC50 values, sensitive (IC50 ≤ maximum testing dosage) and resistant (IC50 > the maximum testing dosage) to calculate prediction accuracy. To measure how precisely the model can predict IC50 values, we calculated MAE for each drug-sample pair belonging to the sensitive class.

#### Combining cell-specific drug response values into an overall response value

IC50 values from CaDRReS-Sc’s pharmacogenomic space provide cell-specific information on the dose-response curve that would have to be integrated across cells to get an overall response profile for a patient. In particular, an average of IC50 values (or weighted average for cell clusters) does not take into account the sigmoid shape of the curve, resulting in inaccurate aggregate IC50 values (naïve estimation, **Supplementary figure 5**). To improve the accuracy of aggregate IC50 calculations, we employed Newton’s method to iteratively approximate the combined dose-response curve based on cell percentages, individual IC50 values and estimated slopes (default=1). The naïve estimate was used to start the iterations, which were observed to converge rapidly in practice.

### Benchmarking drug response predictions for unseen cell types

CaDRReS-Sc was benchmarked against other state-of-the-art machine learning based approaches for drug response prediction, including ElasticNet (Iorio et al., 2016), cwKBMF (Ammad-ud-din et al., 2016), SRMF (Wang et al., 2017), and RWEN (Basu et al., 2018), based on the GDSC dataset and 5-fold cross-validation. For ElasticNet and RWEN, we trained a model for each drug separately based on expression values for all genes. For cwKBMF, we used the same cell line kernel features as CaDRReS-Sc, while excluding drug property information as suggested by the authors. For SRMF, the method does not support prediction for unseen cell lines, as it requires a similarity matrix that consists of both train and test samples. Therefore, we allowed SRMF to use gene expression information for all cell lines but excluded drug response information as appropriate.

### Predicting drug response for head and neck cancer PDCs

Drugs that elicit response in at least 30% (13 out of 42) of head and neck cancer cell lines in the GDSC dataset (n=81) were used to train a head and neck cancer-specific model ( *d*_*u*_ = 10 ; **Supplementary file 4**). The resulting pharmacogenomic space was used to predict cell, cluster and patient-specific drug response values (IC50) based on corresponding transcriptomic profiles. IC50 values were used to estimate cell death percentage for a given dosage *o*_*i*_ and aggregated at the patient level for cell (average) and cell-cluster (weighted average) predictions.

### Drug-pathway associations

A pathway activity score was computed as the summation of gene expression log2 fold-change values across all genes within each BioCarta pathway (Nishimura, 2001). To identify drug-pathway association, the Pearson correlation was calculated between pathway activity scores and predicted drug response values (cell death percentage) across all training samples. Positive correlation coefficient values indicate that high pathway activity is associated with increased drug sensitivity.

### Predicting combinatorial therapy response

To predict combinatorial therapy response, predicted cell death percentages *h*_*i*_ and *h*_*j*_ at specific dosages *o*_*i*_ and *o*_*j*_ of drug *i* and *j* for each cell cluster were aggregated for each cluster as *h*_*i*_ + *h*_*j*_ – *h*_*i*_*h*_*j*_, under the assumption that drugs *i* and *j* inhibit cells independently. To estimate response for a patient, the weighted average of cell death percentages was computed across cell clusters.

The potential utility of a drug combination over individual drugs was calculated as the increase in cell death percentage for the combination compared to the best individual drug within the combination. To prioritize drug combinations for the experimental study, we first confirmed that monotherapy predictions showed high cross-validation accuracy, further identified individual drugs that could inhibit different subclones within a patient, and focussed on combinations that were predicted to improve over monotherapy for at least one patient (**Supplementary file 5**).

### Experimental validation

#### Cell line isolation and cell culture

Cell lines were isolated as mentioned in previous work (Chia et al., 2017). Briefly, tumors were minced and enzymatically dissociated using 4 mg/mL-1 Collagenase type IV (Thermo Fisher, cat. no. 17104019) in DMEM/F12, at 37 °C for 1 hour. Post digestion, cells were pelleted and resuspended in phosphate-buffered saline (Thermo Fisher, cat. no 14190235) for 3 cycles. Cells were then strained through 70 μm cell strainers (Falcon, cat. no. 352350), prior to pelleting and resuspension in RPMI media (Thermo Fisher, cat. no 61870036), containing 10% fetal bovine serum (Gibco, cat. no 10270-106) and 1% penicillin-streptomycin (Thermo Fisher, cat. no. 15140122). Cells were plated on CellBIND plates (Corning, cat. no 3335) and kept in a humidified atmosphere of 5% CO_2_ at 37°C. Cells were routinely screened for mycoplasma contamination using MycoAlertTM PLUS Mycoplasma Detection Kit (Lonza, cat. no: LT07-710).

#### Compounds, drug response and cell viability assays

For each patient, a separate line was isolated from tumor obtained from the patient’s primary and metastatic lymph node sites. Approximately 5000 cells (2500 primary and 2500 metastatic cells) were seeded per well of a 96-well plate, 24 hours prior to drug treatment. Drugs that were used for treatment were obtained from SelleckChem, MedChemExpress and Cayman Chemical. Docetaxel (cat. no. S1148), Doxorubucin hydrochloride (cat. no. S1208), Epothilone B (cat. no. S1364), Obatoclax Meylate (cat. no. S1057), PHA-793887 (cat. no. S1487), PI-103 (cat. no. S1038) and Vorinostat (cat. no. S1047) were obtained from Selleckchem, while Gefitinib was purchased from Cayman Chemical (cat. no. 13166) and Staurosporin (cat. no. HY-15141) from MedChemExpress. Cells were treated at a drug concentration that corresponds to the median IC_50_ value for head and neck cancer cell lines seen in the GDSC database (Iorio et al., 2016), as well as at a concentration that is 3-fold lower (**Supplementary file 6**). All compounds were dissolved in DMSO (Sigma Aldrich, cat. no. D8418) and kept at a constant 1% (v/v) across all drug concentrations and controls. Cells were treated for 72 hours prior to the evaluation of drug response. The amount of viable cells post drug treatment was quantitated using CellTiter-Glo luminescent reagent (Promega, cat. no. G7572). An integration time of 250ms was used when luminescence signals were read using TECAN Infinite M1000 pro multi-mode plate reader. The relative luminescence of each well was computed using the following formula (Luminescence _Drug_/ Luminescence _DMSO_) and expressed as percentage cell viability (**Supplementary file 7**). The median cell death percentage (100 – cell viability) was then calculated across replicates.

### Data availability

The single-cell RNA-seq data used in this study is available in the Gene Expression Omnibus repository under the series GSE117872.

