## Supplementary figures for "Predicting heterogeneity in clone-specific therapeutic vulnerabilities using single-cell transcriptomic signatures"

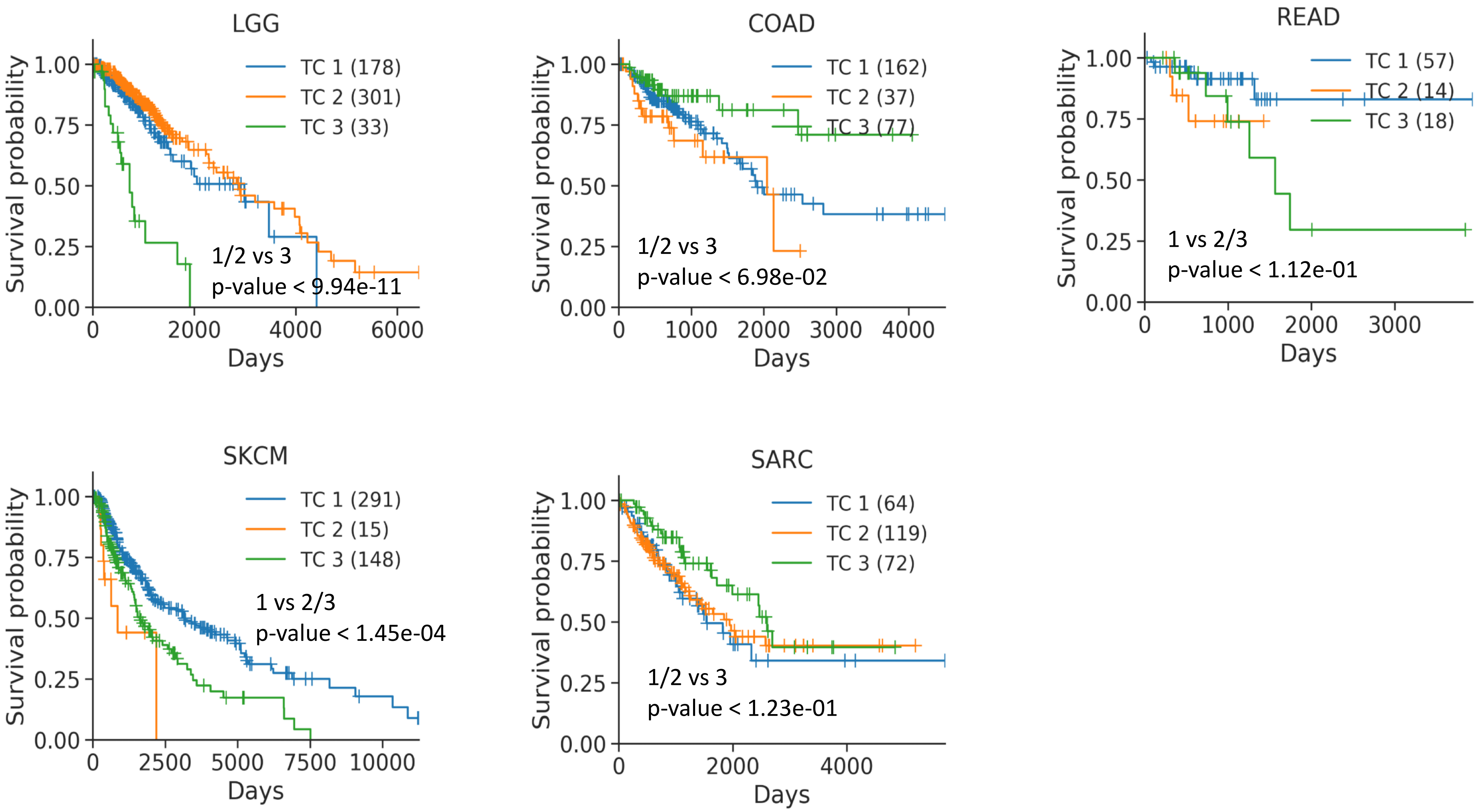


**Supplementary figure 1: Survival analysis for clusters based on bulk transcriptomic profiles.** Note that the figures only show cancer types where significant differences were observed across the clusters. Numbers in parentheses indicate the number of subjects in each cluster. P-values are FDR-corrected and based on the logrank test.

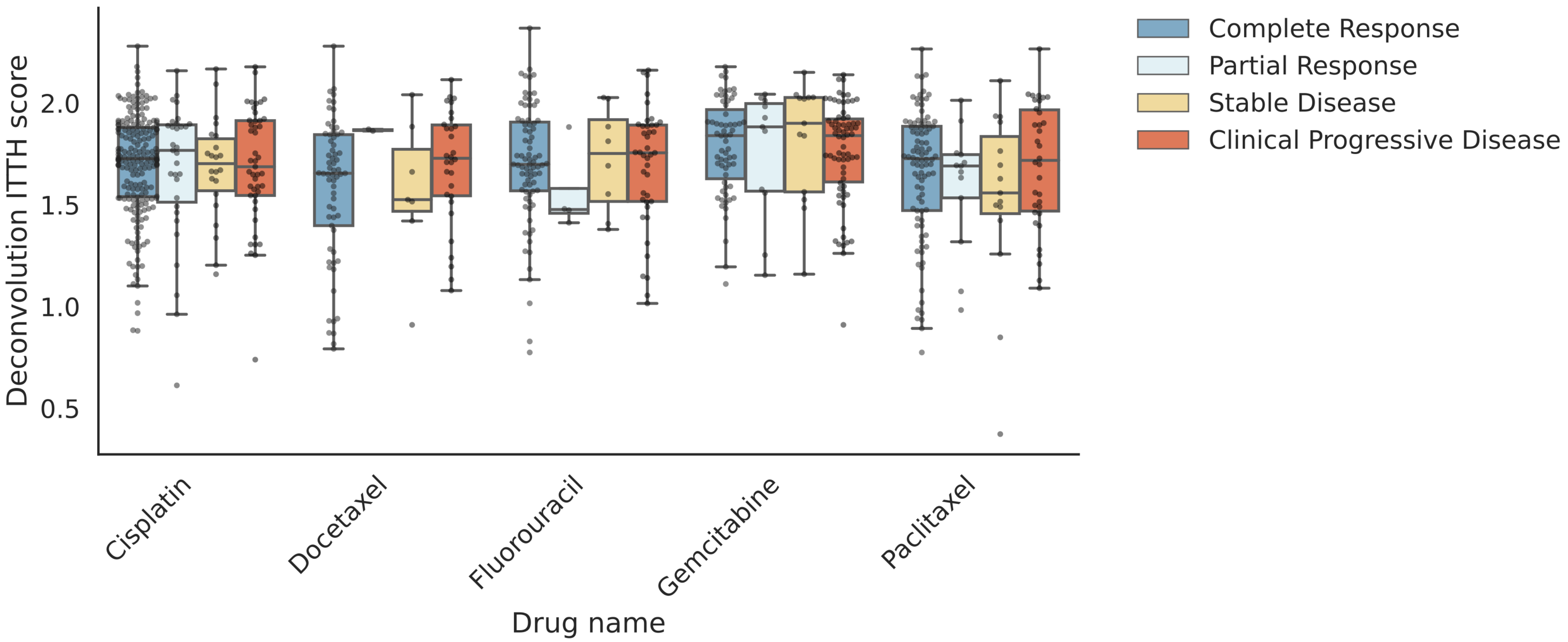


**Supplementary figure 2: Boxplots comparing ITTH scores across clinical response categories for various cancer drugs.** This figure shows drugs that did not exhibit significant differences and the rest are shown in **Figure 1D**.

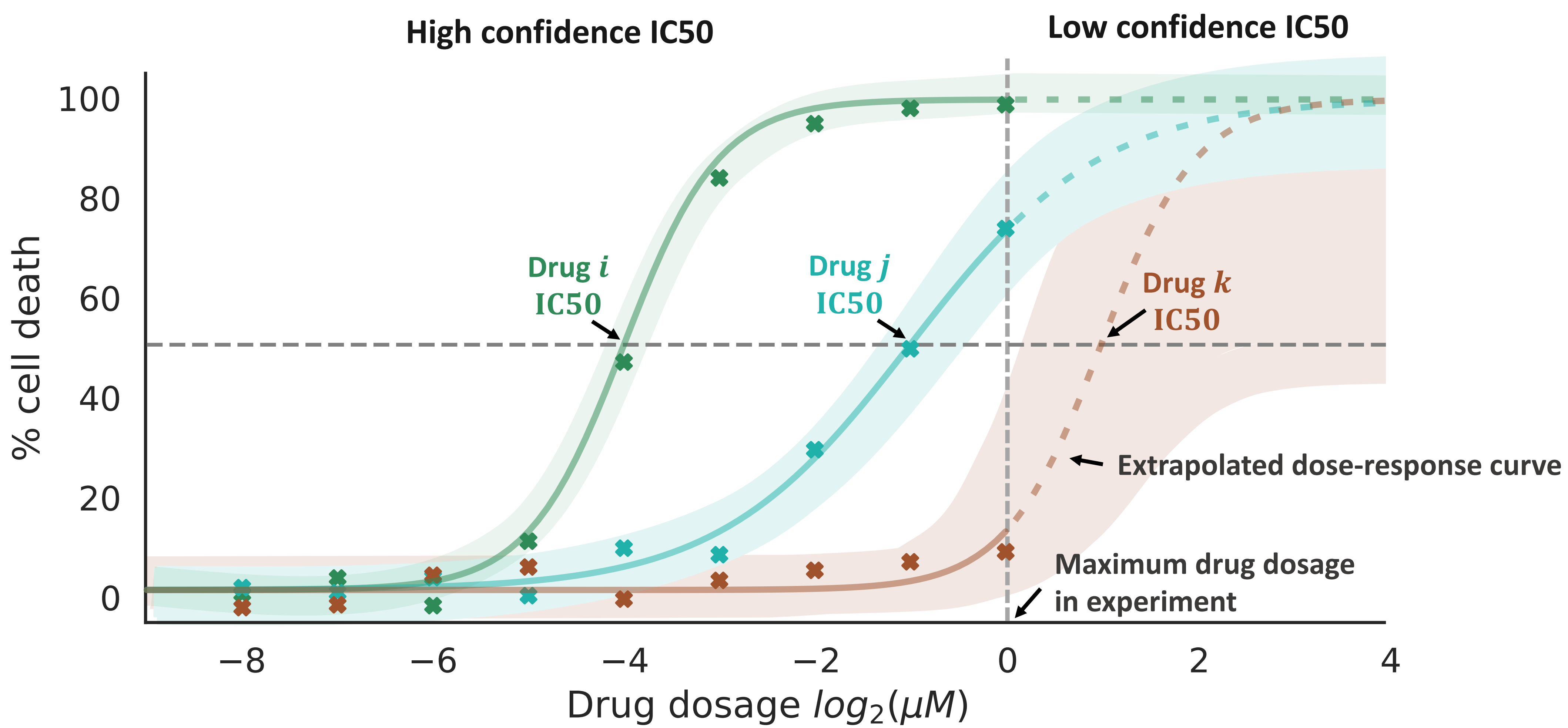


**Supplementary Figure 3: Impact of dose-response curves from *in vitro* cell viability assays on IC50 estimates.** Half-maximal inhibitory concentrations or IC50 values are estimated based on fitting a sigmoid function to the dose-response curve. When true IC50 values are beyond the maximum drug dosage tested, this can lead to significant extrapolation noise that impacts model performance. CaDRReS-Sc addresses this issues with a novel objective function that down-weights low confidence IC50 values during model training (see **Methods**).

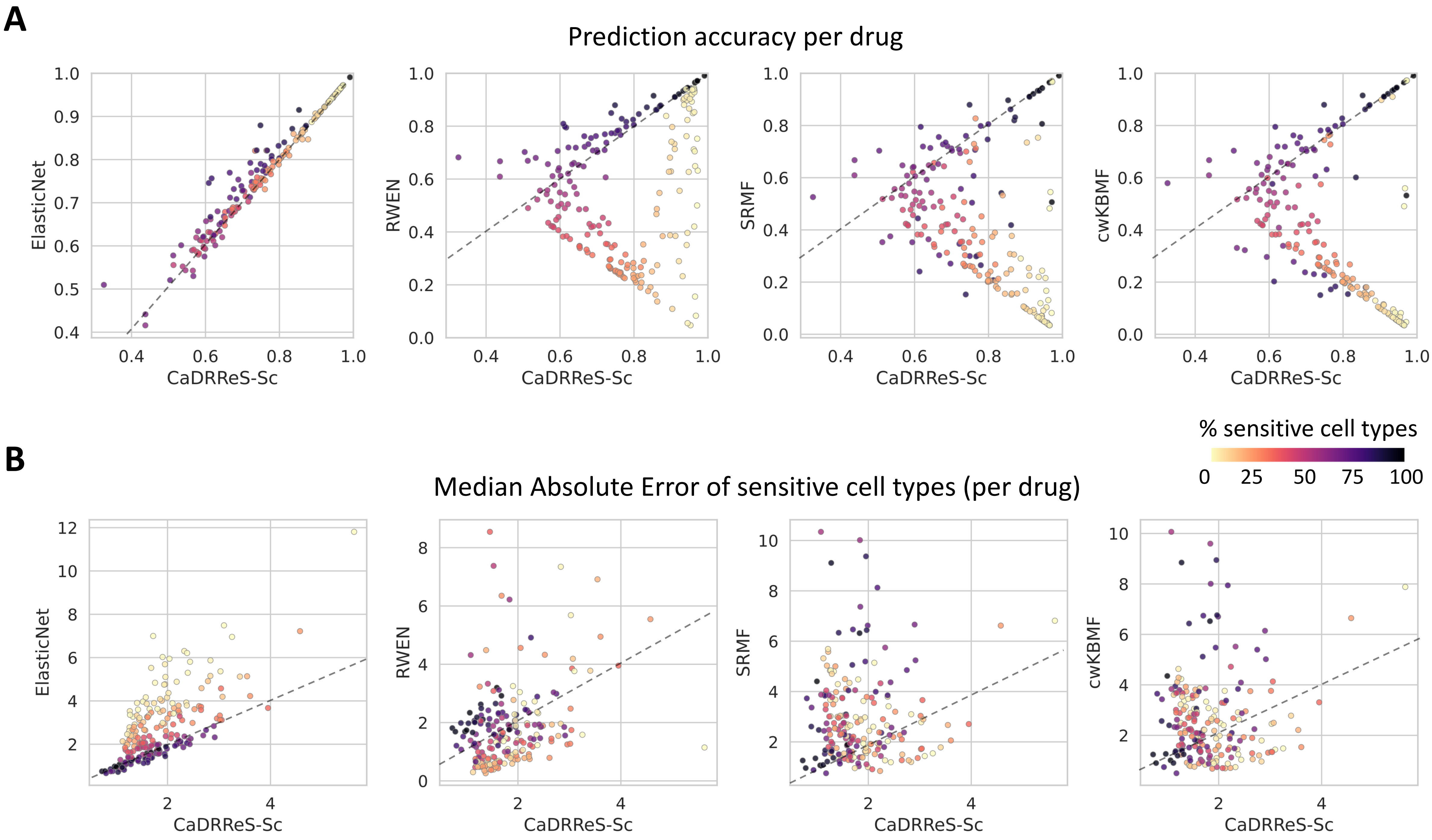


**Supplementary Figure 4: Pairwise comparison of CaDRReS-SC's performance on unseen cell types.** Each dot represents a drug (n=226) and dot colors represent the percentage of sensitive cell types. (A) Comparison of prediction accuracy, where CaDRReS-Sc typically has higher values. (B) Comparison of median absolute error (MAE) on a log2 scale, where CaDRReS-Sc typically has lower values compared to other methods.

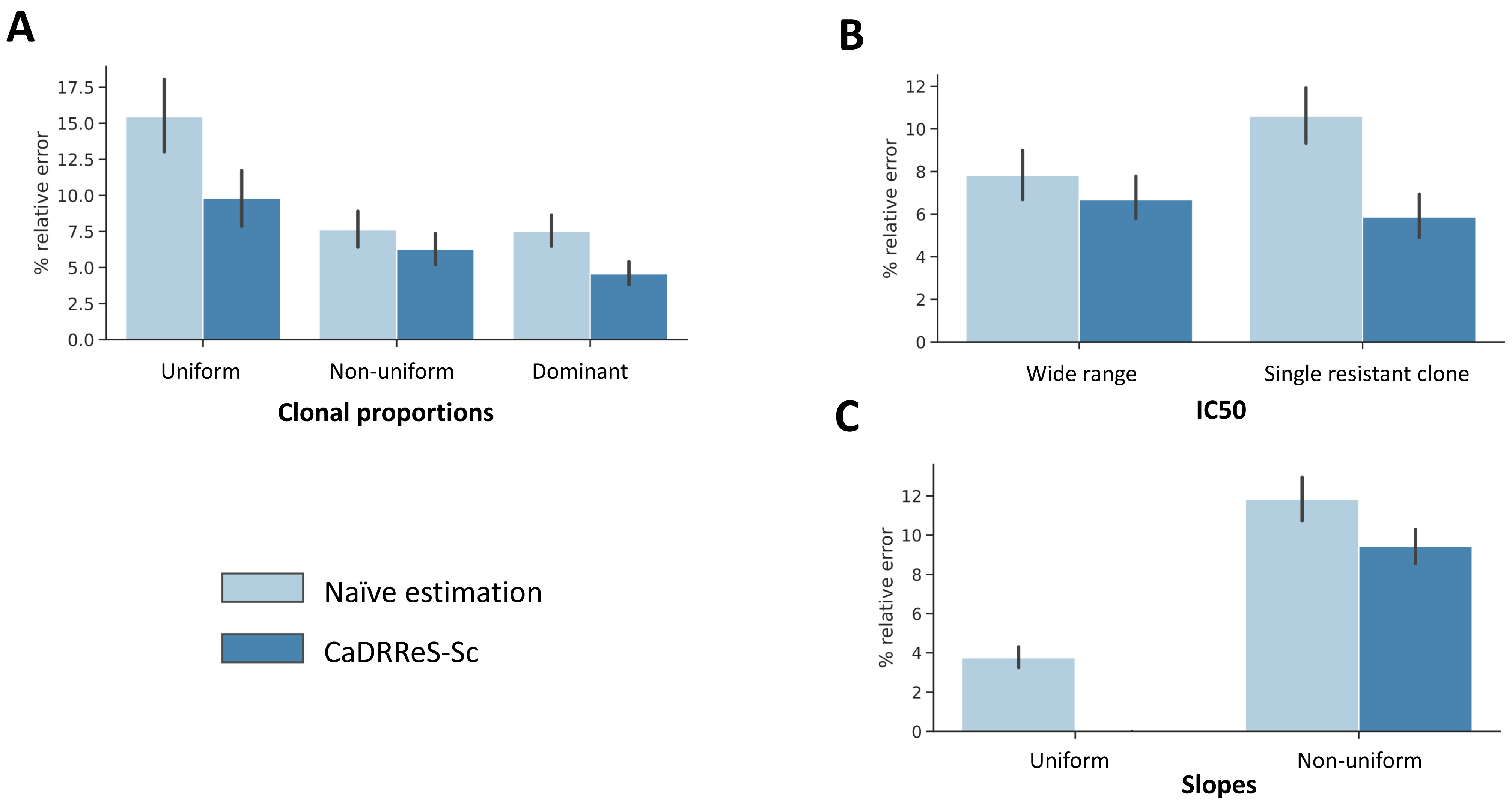


**Supplementary Figure 5: CaDRReS-Sc accurately estimates aggregate IC50 values in the presence of transcriptomic heterogeneity.** Simulated samples were constructed with multiple sub-clones (n=[2-10]) with different dose-response curves based on GDSC data (log2 IC50=[1-5] and slope=[1-3]). CaDRReS-Sc estimates (based on numerical integration; see **Methods**) and a naïve estimation (weighted average of IC50 values) were compared to true IC50 values. Based on different conditions for (A) clonal proportions (B) IC50 values, and (C) slopes, CaDRReS-Sc was observed to be more robust compared to the naïve approach. For clonal proportions, *non-uniform* indicates log distribution abundance and the *dominant* case consist of a single dominant clone (80%) and equal proportions of the remaining clones. For IC50, *wide range* (1-5) indicates different IC50s across clones and *single resistant clone* has a single clone with high IC50 (5) with lower IC50s for the other clones (1). For slopes, *non-uniform* cases refer to the situation with different slopes (1-3) across clones.

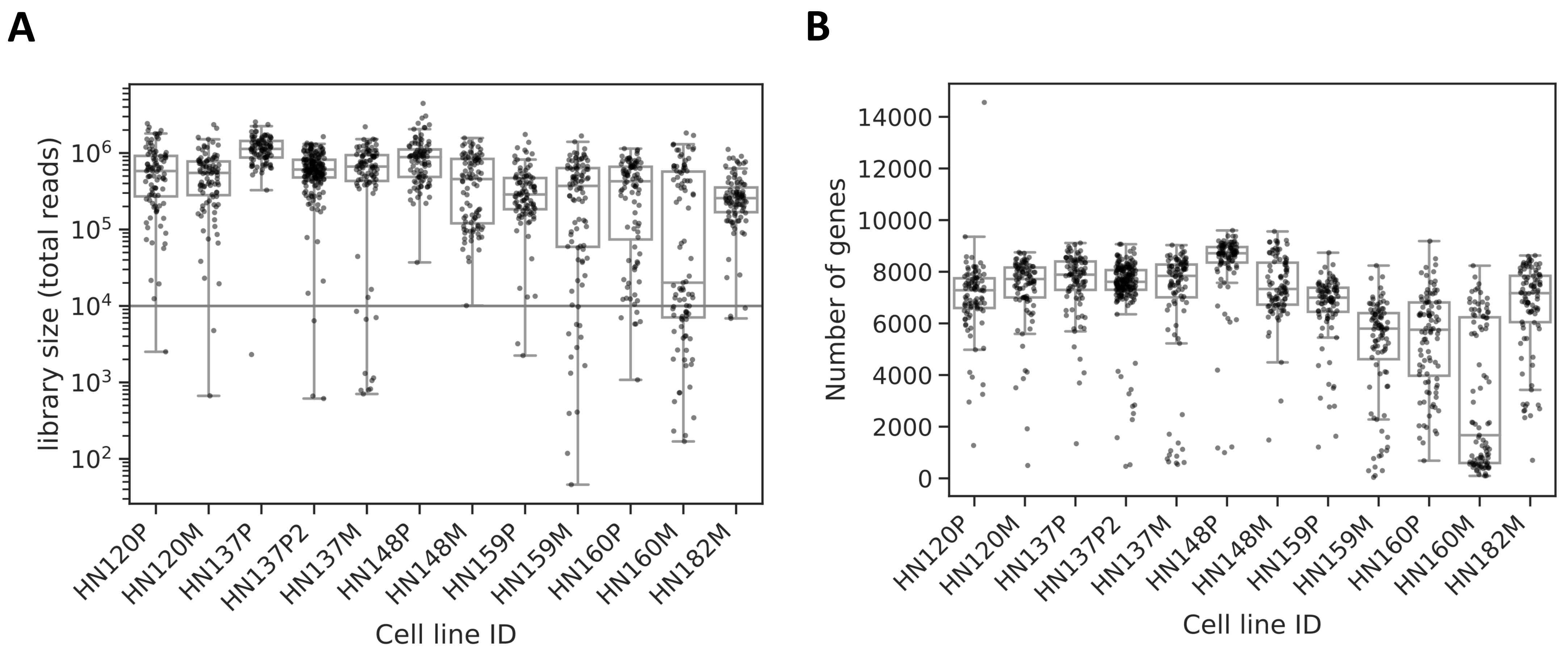


**Supplementary Figure 6: Single-cell RNA-seq statistics for 12 patient-derived cell lines.** (A) Boxplots depicting total number of reads per cell for the 12 cell lines. (B) Boxplots showing total number of expressed genes (read count > 0) per cell for the 12 cell lines.

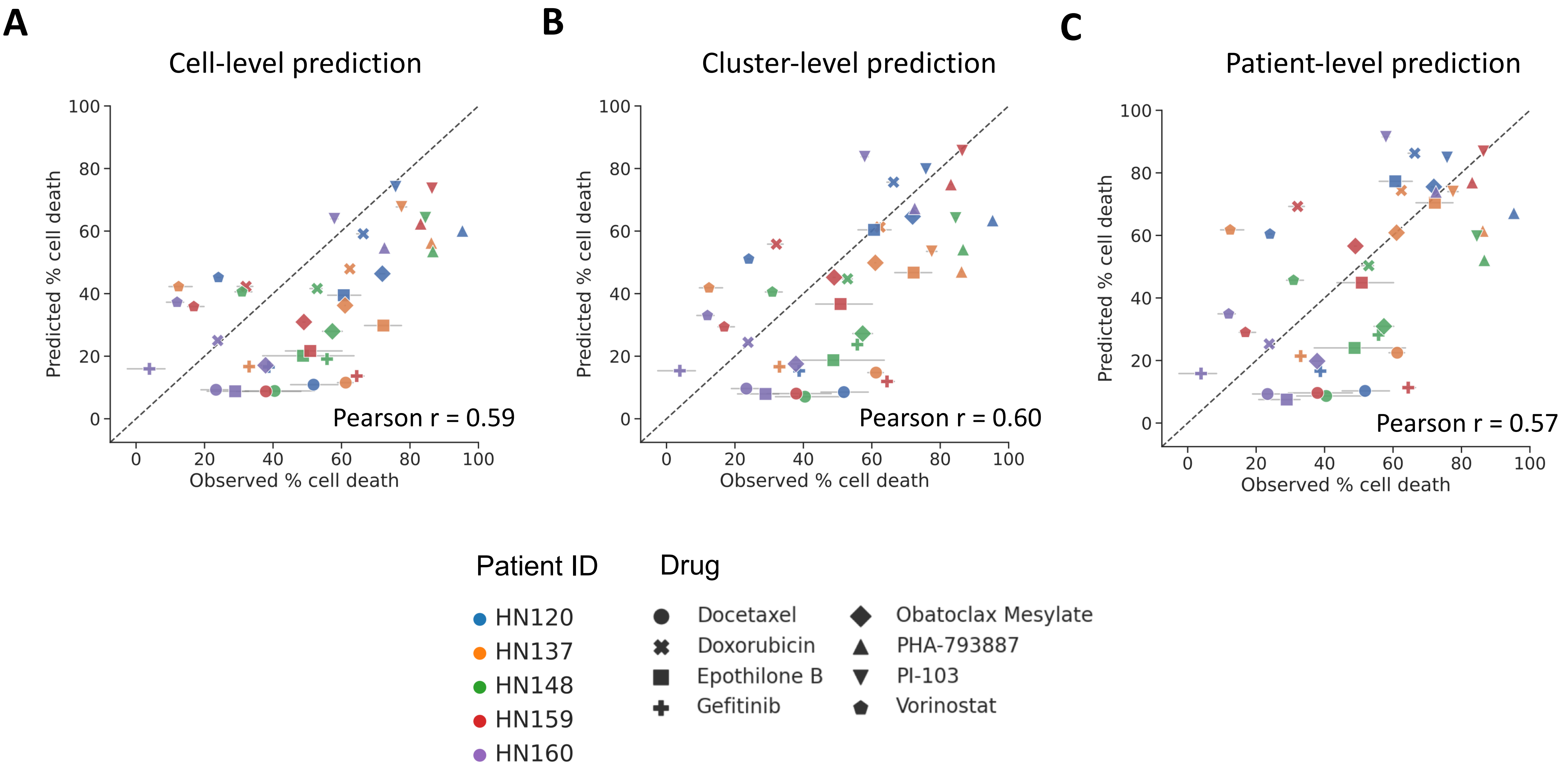


**Supplementary Figure 7: Comparison of observed and predicted drug response across 5 pooled PDCs and 8 drugs.** Prediction based on transcriptomic profiles of (A) each cell combined at the patient-level, (B) each cell cluster combined at the patient-level, and (C) at the patient-level. The results reported here are based on the higher drug concentrations tested.

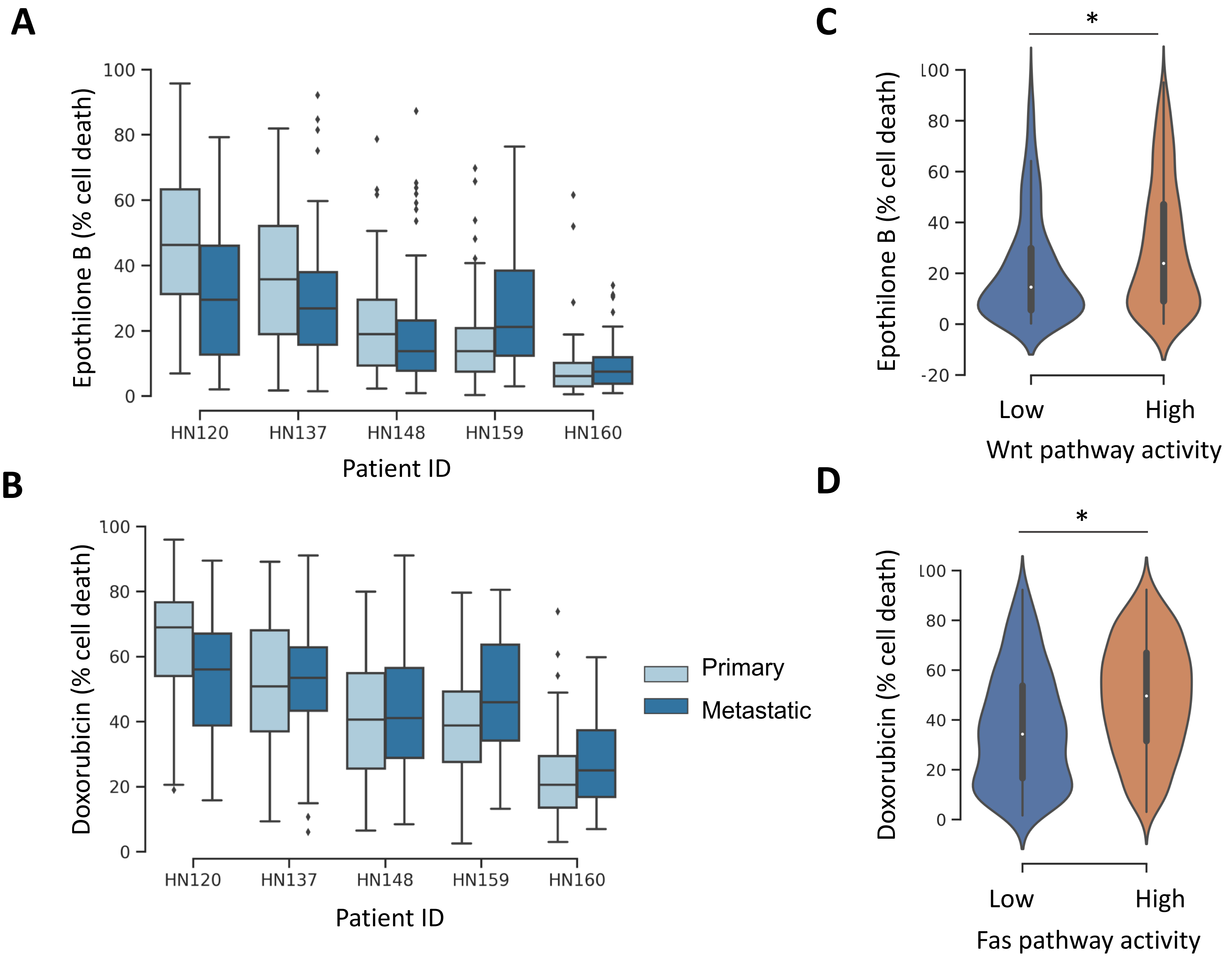


**Supplementary Figure 8**: (A-B) Comparison of predicted drug response from matched primary and metastatic PDCs with CaDRReS-Sc. (C-D) Comparison of pathway activity and predicted drug response in individual cells, highlighting higher sensitivity for cells with higher activity of corresponding pathways (Wnt for Epothilone B and Fas for Doxorubicin).
